# Long-Read Metagenomic Sequencing Reveals Persistent Gut Microbiome Alterations in Children with Celiac Disease in Remission

**DOI:** 10.64898/2026.09.11.749611

**Authors:** David Le, Daniel M. Portik, Takumi Konno, Maureen M. Leonard, Alba Miranda-Ribera, Alessio Fasano, Ali R. Zomorrodi

## Abstract

**Background:** Celiac Disease (CeD) is an immune-mediated enteropathy linked to gut microbiome dysbiosis driven by dietary gluten ingestion. Although most patients achieve clinical and histologic remission on a gluten-free diet (GFD), persistent symptoms and incomplete mucosal recovery are common. While gut microbiome alterations during active CeD are well documented, taxonomic and functional shifts during histologic remission remain considerably less characterized.

**Methods:** We utilized PacBio’s HiFi long-read metagenomic sequencing to profile the gut microbiome of 21 pediatric participants with histologically confirmed CeD remission on a GFD, active CeD, and non-CeD controls.

**Results:** High-fidelity long reads enabled reconstruction of 857 high-quality metagenome-assembled genomes and high-resolution species- and pathway-level profiling. Microbial diversity analysis showed reduced α-diversity in remission patients compared with controls. Differential abundance analysis revealed persistent taxonomic and functional alterations in CeD remission, indicating incomplete restoration of microbiome homeostasis. Notably, remission patients showed depletion of commensals such as *Alistipes finegoldii*, *Bifidobacterium longum*, and *Bacteroides uniformis*, and persistence of some inflammation-associated taxa, alongside enrichment of beneficial species, including short-chain fatty acid producers, and suppression of other inflammation-associated taxa. At the functional level, pathways related to broader metabolic homeostasis were depleted in remission, while anti-inflammatory pathways such as propionate metabolism were enriched.

**Conclusions:** Our findings suggest a distinct microbial ecosystem during histologic remission characterized by residual taxonomic dysbiosis and functional constraints, accompanied by adaptive remodeling that may promote mucosal healing. This work advances understanding of the gut microbiome in CeD after mucosal recovery and lays the groundwork for future studies of microbiome-based therapeutic interventions.

## Background

Celiac Disease (CeD) is a chronic systemic autoimmune condition characterized by enteropathy, which affects over three million Americans and 1.4% of the population worldwide ^1^. In individuals carrying HLA DQ2 or HLA DQ8 alleles, gluten ingestion triggers an immune response leading to enteropathy and subsequent gastrointestinal and extraintestinal symptoms ^2^. However, not all genetically predisposed individuals that are exposed to dietary gluten develop CeD ^1, 3^, indicating that additional factors contribute to its pathogenesis. Among these, the gut microbiota has recently emerged as a key contributor ^4-7^.

The microbiome may contribute to CeD pathogenesis through several mechanisms, including modulation of gluten peptide immunogenicity, intestinal barrier integrity, and mucosal immune tolerance ^8^. Previous studies of both duodenal and fecal microbiota have indicated that CeD is characterized by a decrease in abundance of beneficial species, such as *Lactobacillus* and *Bifidobacterium*, with concurrent enrichment of pathogenic species, such as species of *Bacteroides*, *Escherichia,* and *Neisseria* ^4, 6^. Analysis of the stool microbiome in at-risk infants using a prospective birth cohort study in our prior work further demonstrated that microbial shifts could precede disease onset, suggesting a possible causal role for dysbiosis in initiating or amplifying intestinal inflammation in CeD. ^9, 10^. Notable alterations included the increased abundance of several gut microbiome features linked to inflammatory conditions (e.g., *Dialister invisus*, *Lachnospiraceae bacterium*, tryptophan metabolism, and the metabolites serine and threonine) and reduced abundance of anti-inflammatory features (e.g., *Streptococcus* thermophilus, *Faecalibacterium prausnitzii*, and *Clostridium clostridioforme*) prior to CeD onset ^10^.

Despite the accumulating evidence, the therapeutic potential of microbiome-targeted interventions in CeD has not been fully explored, and strict adherence to a gluten-free diet (GFD) remains the only established treatment ^11^. However, some patients continue to experience persistent symptoms or incomplete mucosal recovery despite long-term dietary restriction. This suggests that GFD treatment may not fully restore intestinal homeostasis in patients, pointing to the potential involvement of the gut microbiome in achieving and/or maintaining clinical and histologic remission. Nevertheless, while the gut microbiome dysbiosis in active CeD has been investigated extensively and is relatively well characterized, microbiome shifts during histologic remission are underexplored and it remains unclear whether the gut microbiome normalizes in patients on a GFD with confirmed mucosal recovery. Research on other conditions, such as inflammatory bowel disease (IBD), has shown that dysbiosis persists even during remission ^12, 13^, suggesting that similar trends might exist for CeD remission.

Prior research examining gut microbiome alterations in GFD-treated CeD patients ^14-17^, has provided important evidence of persistent alterations; however, the evidence remains limited and methodologically heterogeneous. Importantly, many of these studies did not include contemporaneous assessment of mucosal histology and therefore could not establish whether participants had achieved mucosal recovery, leaving the microbiome during histologic remission incompletely characterized. Zafeiropoulou et al ^14^ and Sample et al ^15^ both used 16S rRNA amplicon sequencing to profile the gut microbiota of pediatric CeD patients on a GFD. These studies identified microbial shifts associated with a GFD— such as increased abundance of taxa like *Clostridium sensu* stricto 1 and *Ruminococcus* ^14^ and reductions in fiber-degrading taxa like *Blautia*, *Dorea*, and *Lactobacillus* ^15^—most of which were attributed to the influence of GFD rather than disease activity alone. However, the reliance of these studies on amplicon sequencing limited resolution to higher taxonomic levels (like genus and above) and precluded functional analysis. In contrast, Francavilla et al. ^16^ employed shotgun metagenomics alongside small RNA profiling in adult CeD patients and found that even those treated with a strict GFD who had negative serology exhibited persistent microbiome alterations. These included alterations in both species composition (e.g., reduced abundance of *Bifidobacterium longum* and *Ruminococcus bicirculans*) and microbial pathways (e.g., altered pathways related to denitrification). More recently, the microbiome analysis of duodenal aspirates found impaired microbial fiber metabolism and depletion of fiber-degrading *Prevotella* spp. in both active and GFD-treated CeD ^18^. Metabolomic evidence also suggests incomplete normalization after treatment, with several fecal metabolites remaining altered in GFD-treated pediatric CeD patients relative to healthy controls ^19^. Collectively, these studies suggest that the gut microbiome does not fully normalize following GFD treatment; however, the taxonomic and functional features associated with histologic remission remain poorly resolved, as there is currently very limited evidence evaluating the microbiota of patients with documented mucosal recovery.

Recent advancements in long-read sequencing technologies have transformed microbiome research by offering greater read lengths—e.g., 10 to 20 kilobases for Pacific Biosciences’ (PacBio’s) HiFi sequencing—while maintaining comparable base calling accuracy ^20^. These longer reads capture repetitive and structurally complex genomic regions that short-read methods often fail to resolve, allowing for the identification of full-length genes, operons, and other genomic features with high precision. These features enable more accurate resolution of species, subspecies, as well as functional profiles and facilitate the assembly of high-quality and near-complete metagenome-assembled genomes (MAGs). Long-read MAGs allow for the detection of novel microbial species and provide direct access to biosynthetic pathways, virulence genes, and metabolic capacities that are often fragmented or overlooked in short-read assemblies.

In this study, we leveraged long-read metagenomic sequencing to investigate stool microbiome alterations in pediatric patients with CeD on a GFD in histologic remission compared to those with active CeD and non-CeD controls both on a regular gluten containing diet. By combining read-based profiling with genome-resolved MAG analysis, we sought to define species- and functional-level microbial changes associated with remission and provide a deeper genome-informed view of the microbiome landscape that may contribute to sustaining disease recovery.

## Results

### Participant cohort and sequencing overview

The study cohort comprised 21 pediatric participants (52.38% female; mean age: 10.38 years; age range: 3–20 years) recruited at Massachusetts General Hospital, Department of Pediatrics (Boston, MA). The cohort includes 6 non-CeD controls, 9 individuals with active CeD, and 6 patients with CeD in histologic remission (CeD remission; Marsh 0–1) following a GFD (mean duration on GFD: 2.8 years; range: 1–6 years) (see **Supplementary File 1**). Stool samples from all 21 participants underwent long-read metagenomic sequencing using PacBio’s HiFi technology. These high-fidelity sequencing reads were used for MAG reconstruction, taxonomic profiling, and functional profiling.

### MAG assembly

Assembly of HiFi reads yielded a total of 857 high-quality MAGs, each with <20 contigs, <10% contamination, and >70% completeness (**Figure 1A**). These MAGs ranged in size from 1 to 7 Mbp and could be assembled with as little as 5X depth of coverage (**Figure 1B**). Notably, 337 MAGs (39.3%) were of new-complete quality—defined by a single contig, >95% completeness, and <5% contamination—demonstrating the ability of long-read HiFi sequencing to reconstruct near-complete microbial genomes. On average, each sample contributed 41 MAGs, of which 16 were classified as near-perfect quality (**Figure 1C**).

**Figure 1.**
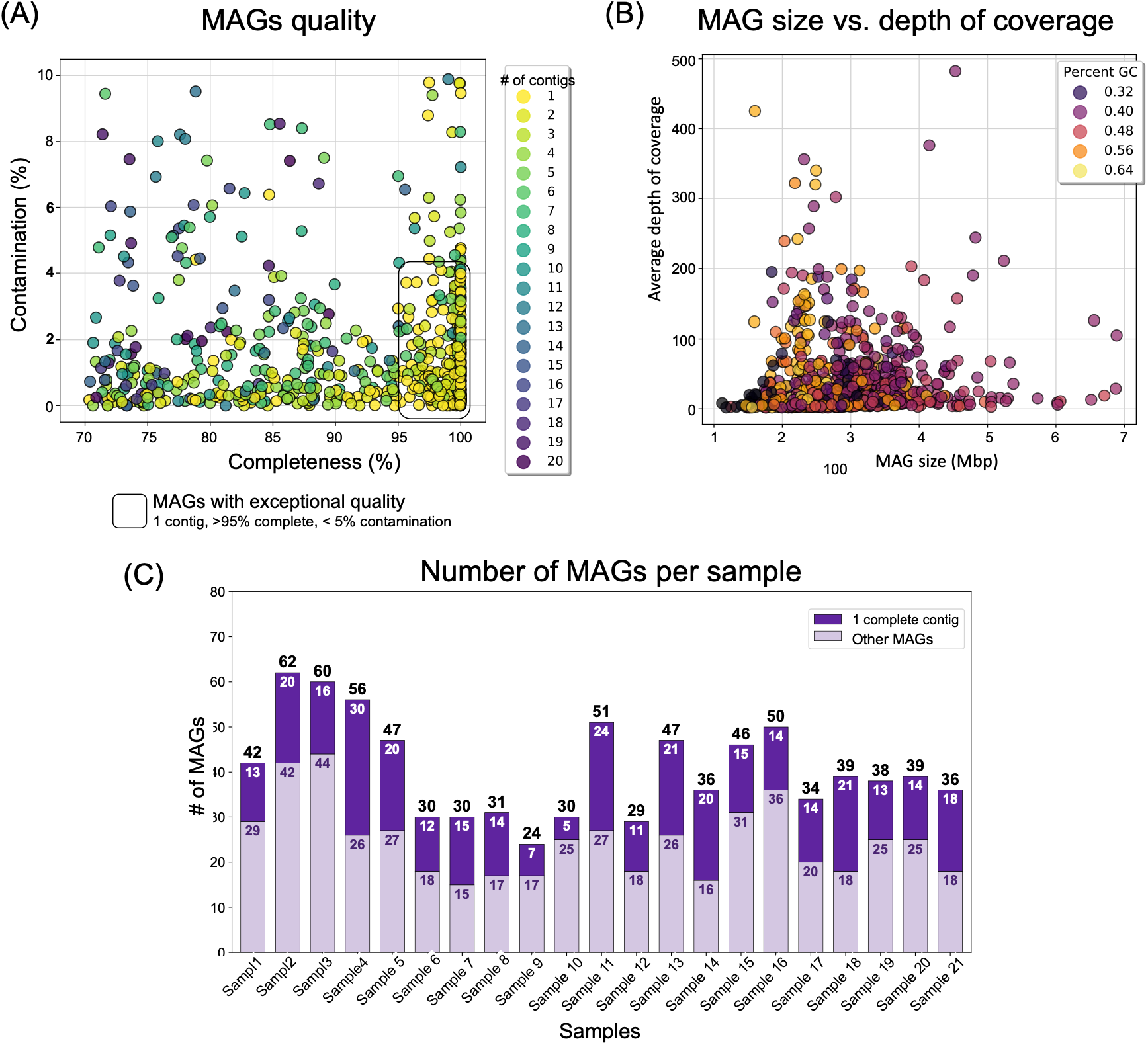
Assembly and quality assessment of MAGs. (A) Completeness vs. contamination for the 857 high-quality MAGs assembled from HiFi long-read reads. (B) MAG size vs. depth of coverage. (C) The number of MAGs recovered per sample, including the subset of exceptional-quality MAGs defined by single-contig assemblies, >95% completeness, and <5% contamination.

### Taxonomic and functional profiling

Taxonomic profiling was performed using both long-read alignment and MAG-based approaches. Read-based profiling identified 1,361 distinct microbial species across the 21 samples, while MAG-based profiling detected 251 unique species across the 836 MAGs. Notably, 21 MAGs lacked annotation in current reference databases and could not be assigned to any known species. This highlights the ability of long-read genome reconstruction to capture previously uncharacterized microbial genomic diversity by identifying potentially novel microbial species that may not yet be represented by cultured isolates or reference genomes. Functional profiling based on KEGG Orthology (KO) using read alignment identified 205 distinct metabolic pathways. To ensure robustness in downstream analyses, both taxonomic profiling results from read alignment and functional profiles were filtered using a dual threshold: species or pathways were retained only if their relative abundance exceeded 0.1% in at least 10% of samples. After applying these filters, a total of 367 microbial species and 90 pathways were retained for downstream analyses. All the taxonomic and functional profiling results are provided in **Supplementary File 1**.

### Diversity analysis

Alpha diversity was assessed using three metrics: Chao1, Shannon index, and Faith’s Phylogenetic Diversity. Across all three measures, the CeD remission group demonstrated a reduction in microbial diversity compared to both active CeD and control groups (**Figure 2A** and **2B**). Although these differences did not reach statistical significance (Wilcoxon Rank Sum [Mann–Whitney U] test, p < 0.05) for both read-aligned and MAG-derived taxonomic profiles, the pattern suggests constrained microbial diversity during remission. We additionally performed alpha diversity analysis on KEGG pathway profiles based on Chao1 estimator and Shannon index; however, this analysis showed no significant differences (Wilcoxon Rank Sum test, p < 0.05) or a clear trend across the groups (**Supplementary Figure 1A**). Beta diversity analysis using Bray–Curtis dissimilarity, UniFrac ^21^, and weighted UniFrac ^22^ also revealed no significant global community differences among the study groups (PERMANOVA, p < 0.05, **Supplementary Figure 1B and 1C**).

**Figure 2.**
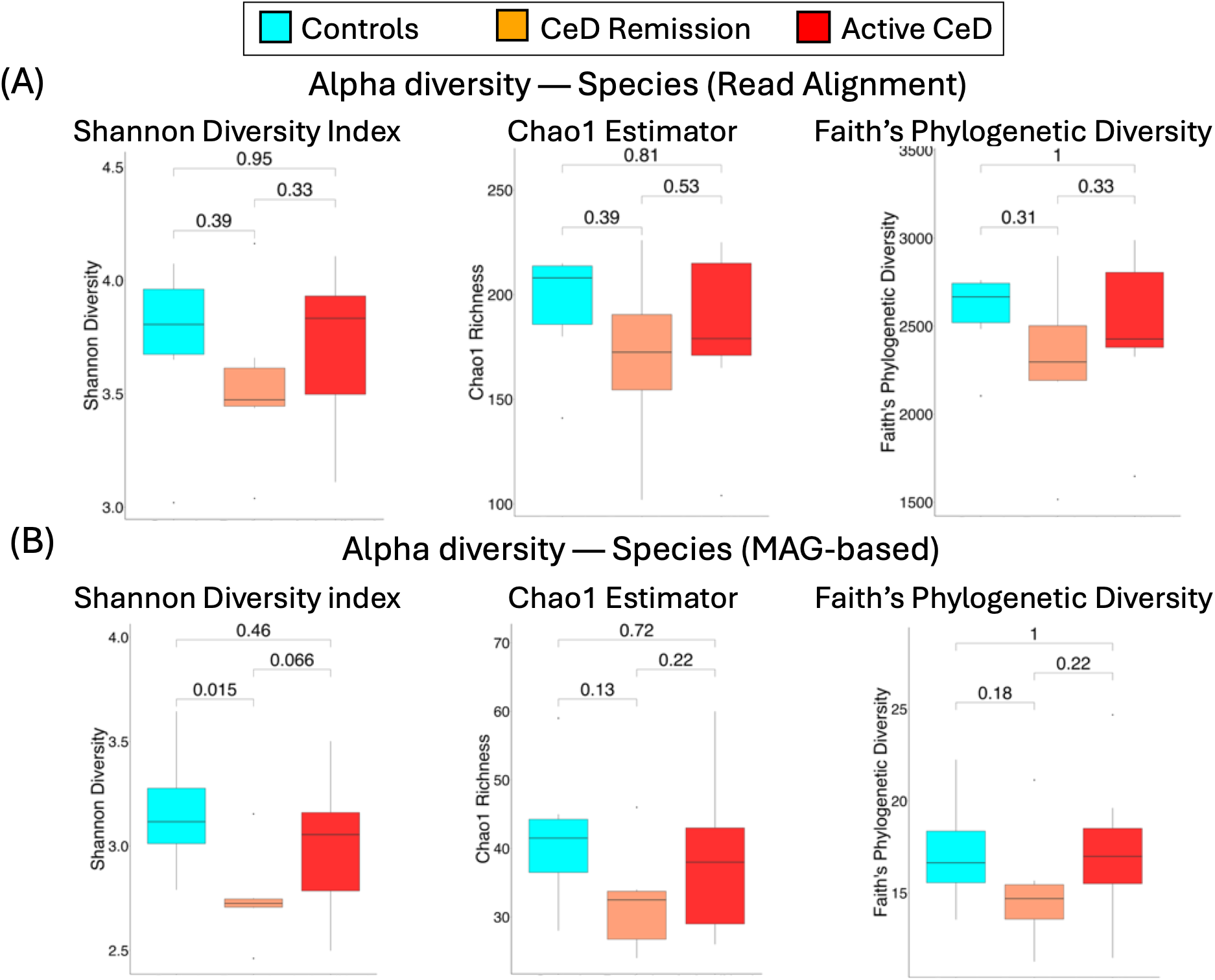
Alpha diversity of gut microbiomes among CeD patient groups and controls. (A) Alpha diversity metrics based on species identified through read alignment, and (B) species assignments from MAGs comparing CeD remission, active CeD, and non-CeD control groups.

### Differential abundance analysis of microbial taxa and pathways

Differential Abundance (DA) analysis was performed to determine microbial species and pathways exhibiting significantly altered abundance between the study groups. Taxonomic DA analysis was performed on microbial species identified through both read alignment and MAG-based approaches, while functional DA analysis focused on KEGG pathways. We conducted three pairwise comparisons: CeD remission vs. active CeD, CeD remission vs. controls, and active CeD vs. controls, using ANCOM-BC ^23^. However, since ANCOM-BC is known to be a somewhat conservative method, we complemented this analysis with a Wilcoxon Rank Sum (Mann-Whitney U) test on the Trimmed Mean of M-values (TMM)-transformed count data. The final DA results were derived by reconciling findings from both methods. Statistical significance was determined using a threshold of p < 0.05 for either ANCOM-BC or Wilcoxon tests. All raw p-values and adjusted p-values for multiple comparisons, for both species and pathways, are provided in **Supplementary File 2**.

#### Differentially abundant species identified via read alignment

A total of 23 microbial species identified through read alignment were determined to be differentially abundant across at least one of the three pairwise comparisons (ANCOM-BC or Wilcoxon; p < 0.05; **Figure 3**). This includes species that were enriched only in CeD remission compared to controls (e.g., *Lachnospiraceae bacterium* 210521-DFI.1.109 and *Ruminoccocus lactaris),* only in active CeD compared to controls (e.g., *Bacteroides fragilis*, *Bacteroides xylanisolvens*, *Faecalibacterium sp.* Marseille-Q4137), or in both CeD remission and active CeD compared to the control group (e.g., *Mogibacterium sp.* NSJ-24). We also identified species that were depleted in CeD remission (e.g., *Eggerthellaceae bacterium*, *Lacrimispora saccharolytica*) or active CeD (e.g*., Faecalibacterium sp.* CLA-AA-H233, *Gordonibacter urolithinfaciens*) compared to controls. Given the low sample size and the associated risk of missing true effects (false negatives), we additionally reported, for descriptive purposes only, the top two microbial species in each pairwise comparison that, while not reaching statistical significance (p < 0.05) in either ANCOM-BC or Wilcoxon tests, exhibited the highest fold changes in TMM-normalized counts in each pairwise comparison. This resulted in an additional five unique microbial species, namely *Blautia caecimuris*, *Blautia hydrogenotrophica*, *Blautia schinkii*, *Blautia sp* 210820-DFI.6.14, and *Dorea sp.* AM13-35 (**Figure 3**).

**Figure 3.**
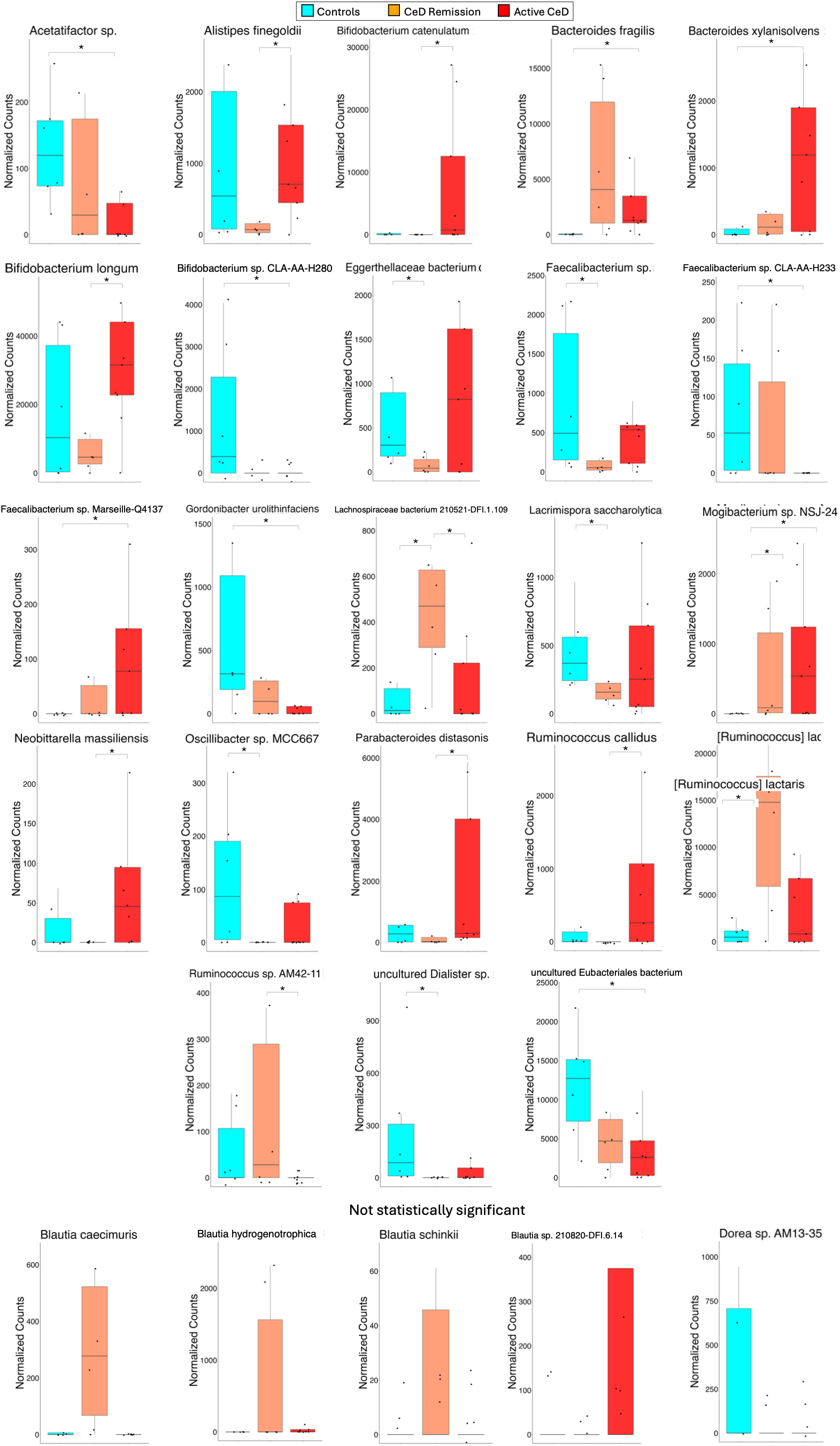
Differentially abundant species identified through read alignment among CeD patient groups. Significant results are based on either ANCOM-BC or the Wilcoxon Rank Sum (Mann-Whitney U) test (p < 0.05) for at least one pairwise comparison among the study groups: active CeD vs. controls, active CeD vs. CeD remission, and CeD remission vs. controls. To mitigate potential false negatives, the top two species with the highest fold change in each pairwise comparison are also shown, resulting in five additional distinct species. The vertical axis indicates TMM-normalized counts. All outliers were removed for visualization purposes, but retained in DA analysis. Outliers were defined using Tukey’s rule (values greater than Q3 + 1.25 × IQR within each group) ^24^. All p-values and corresponding q-values are provided in **Supplementary File 2**.

#### Differentially abundant species identified via MAG-based taxonomic profiling

For MAG-based DA analyses, ANCOM-BC was applied to average depth of coverage values, while Wilcoxon testing was performed on CLR-transformed relative abundances derived from those values (see **Methods** for details). Six microbial species assigned to MAGs showed significantly altered abundance between the study groups (ANCOM-BC or Wilcoxon, p < 0.05) (**Figure 4**). Among these, *Bifidobacterium longum* was depleted in CeD remission compared to active CeD, while *Eggerthella lenta* was enriched in CeD remission compared to both controls and active CeD. Conversely, *Dorea_A longicatena* was depleted in active CeD compared to controls, while three species were enriched in active CeD compared to controls (e.g. *Oliverpabstia intestinalis*, *Romboutsia timonensis*), or both controls and CeD remission (e.g. *Fimenecus sp000432435*). Similar to our DA analysis for species identified through read alignment, we additionally reported the top two non-significant microbial species with the highest fold change in abundance for each pairwise comparison for exploratory purposes. This secondary analysis identified four unique species, which included *Anaerostipes hadrus_A, Bacteroides uniformis, CAG-103 sp000432375,* and *Coprococcus eutactus_A*. Notably, of the reported species shown in **Figure 4**, *B. longum* was identified as differentially abundant in DA analysis of species identified via read alignment-based taxonomic profiling (**Figure 3**) and shows a consistent pattern of depletion in CeD remission in both **Figures 3** and **4**.

**Figure 4.**
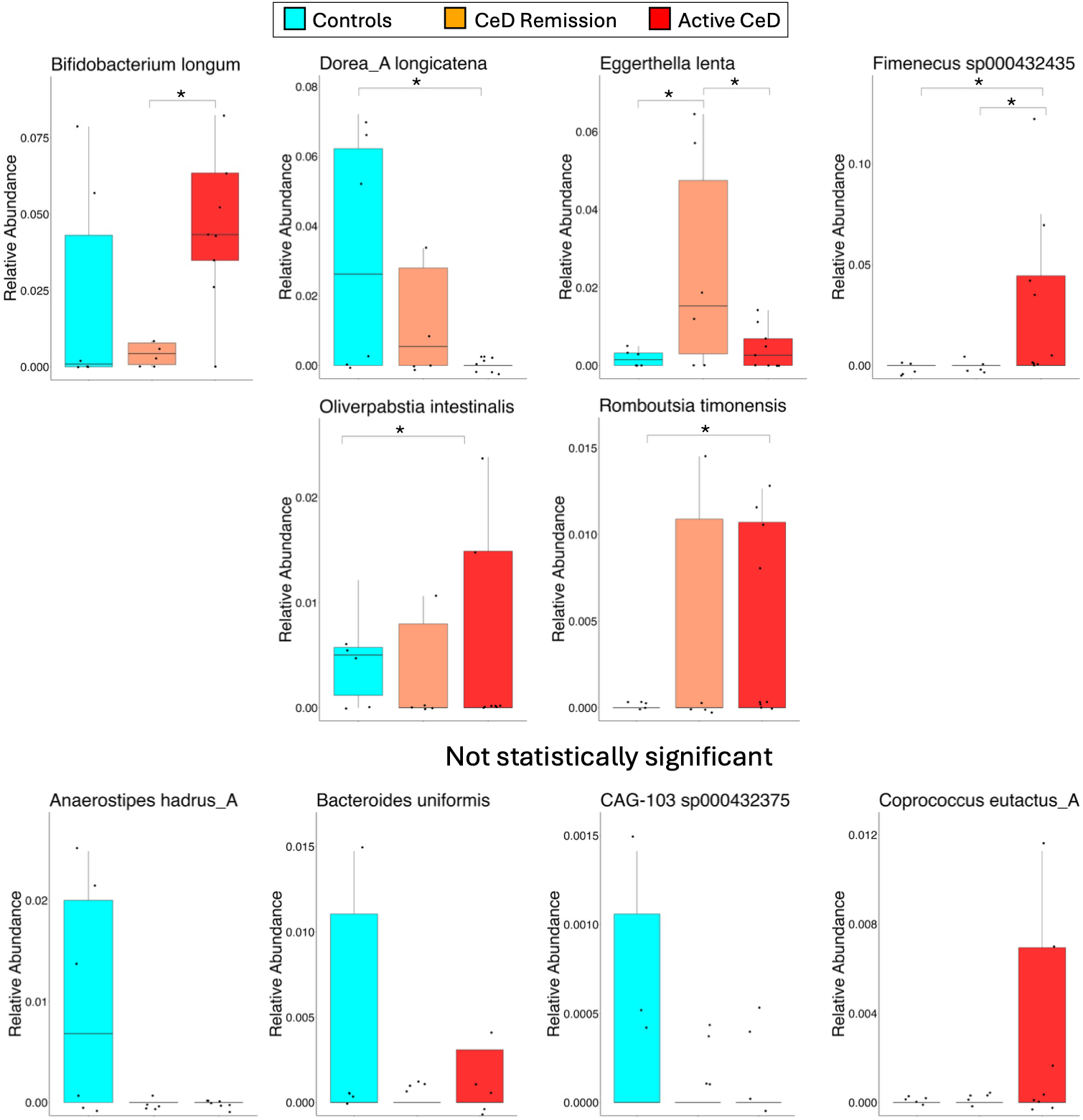
Differentially abundant species identified through MAG-based profiling among CeD patient groups and controls. Statistical significance was determined as described in Figure 3. The top two species exhibiting the highest fold change in each pairwise comparison that did not reach statistical significance (ANCOM-BC or Wilcoxon, p < 0.05) are also presented, resulting in four additional species. The vertical axis indicates relative abundance, calculated from MAG coverage as the average depth of coverage of a species divided by the total average depth across all MAG-derived species within the same sample. Outlier handling was performed as described for Figure 3. All p-values and corresponding q-values are provided in **Supplementary File 2**.

To provide a clearer visualization of microbial shifts associated with each disease state, we consolidated the differentially abundant species shown in **Figures 3** (read alignment-based) and **4** (MAG-based) and reorganized them according to their enrichment or depletion in each disease state (active or remission), which are presented in **Supplementary Figures 2 and 3**.

#### Differentially abundant functional pathways

DA analysis of KEGG pathways identified four pathways exhibiting significantly altered abundance between at least one pair of the study groups (ANCOM-BC or Wilcoxon, p < 0.05) (**Figure 5**). In particular, pentose and glucuronate interconversions was depleted in CeD remission compared to controls while pyrimidine metabolism is depleted in CeD remission compared to active CeD. Conversely, benzoate degradation and propionate (propanoate) metabolism are enriched in CeD remission compared to active CeD. Reporting the top two pathways exhibiting the highest fold change in abundance for each pairwise comparison that did not meet the significance threshold (p < 0.05) to safeguard against potential false negatives highlighted two additional pathways—fatty acid biosynthesis and nitrotoluene degradation—which are enriched and depleted in CeD remission, respectively.

**Figure 5.**
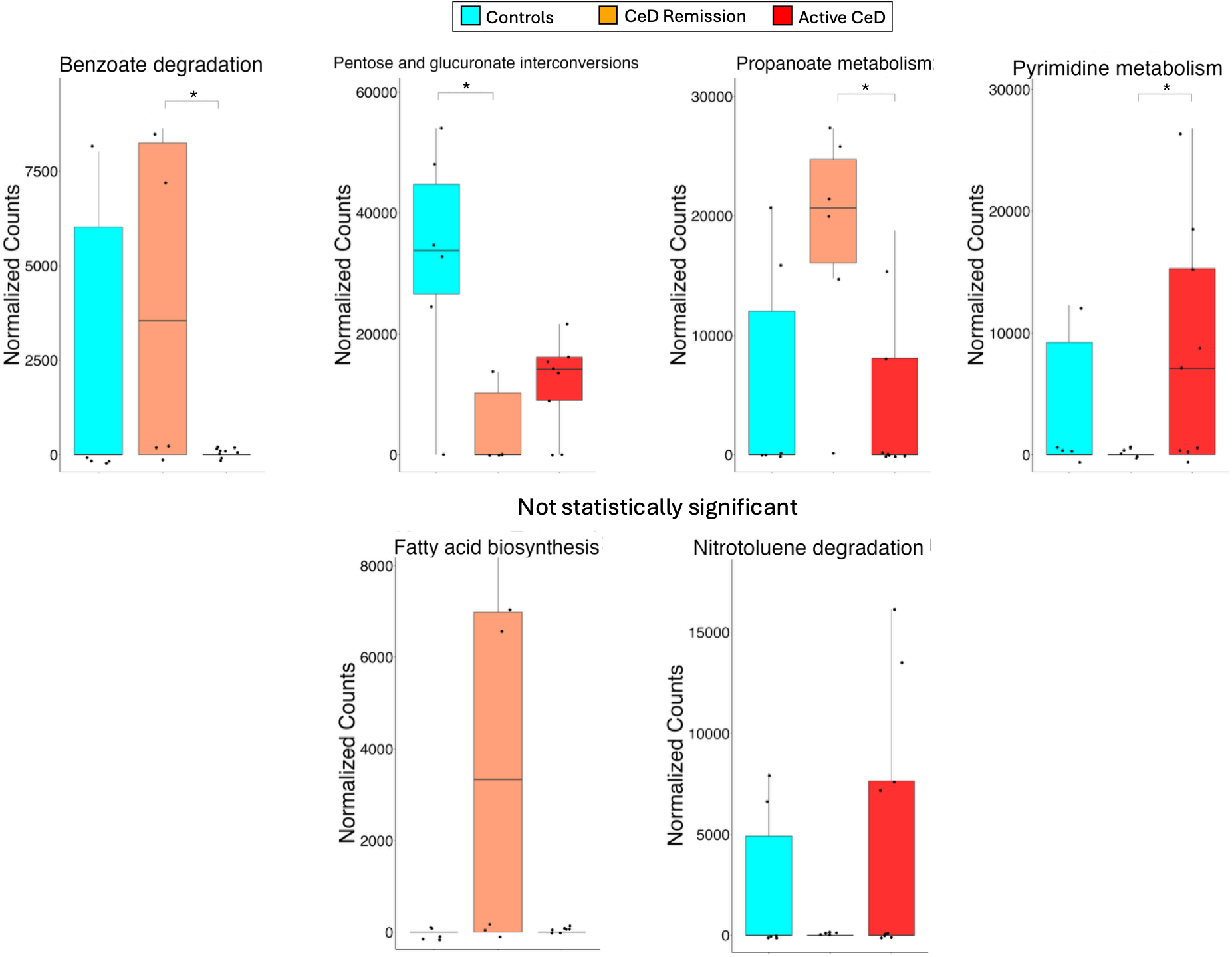
Differentially abundant functional pathways among CeD patient groups and controls. Statistical significance was determined as described in Figure 3. The top two pathways with the greatest abundance fold change in each comparison that did not reach statistical significance (ANCOM-BC or Wilcoxon, p < 0.05) are also presented, resulting in two additional pathways. The vertical axis indicates TMM-normalized read counts. Outlier handling was performed as described for Figure 3. All p-values and corresponding q-values are provided in **Supplementary File 2**.

## Discussion

This study provides a high-resolution characterization of the gut microbiome taxonomic and functional alterations in pediatric patients with CeD during histologic remission, extending prior studies of patients with CeD on a GFD through leveraging long read metagenomic sequencing. Specifically, we sought to determine whether mucosal recovery is accompanied by restoration of a healthy gut microbiome or whether it reflects a distinct equilibrium shaped by dietary restriction and residual immune activation. The use of long-read sequencing enabled reconstruction of 857 high-quality MAGs, including hundreds of near-complete single-contig genomes. Furthermore, because individual long reads generated for our samples were up to 15 Kbp in length, they could span complete genes and operons, providing substantially richer sequence context for taxonomic assignment and especially for functional annotation versus short reads. The integration of reference-based profiling with high-quality MAG-based analysis provided complementary views of community composition and supported a more genome-informed view of remission-associated microbiome shifts. Microbial species and pathways identified through taxonomic and functional profiling of long reads and MAGs were subjected to alpha and beta diversity analysis as well as DA analysis to evaluate gut microbiome alterations in CeD remission relative to active CeD and controls. Our finding of persistent microbiome alterations despite mucosal recovery is broadly consistent with prior studies of GFD-treated CeD ^14-17^; however, our work extends these studies by specifically examining the microbiome in patients with documented histologic remission, a state that has been largely uncharacterized in prior reports.

### CeD remission shows a trend toward reduced alpha diversity following long-term adherence to a gluten-free diet

While no statistically significant changes were observed between the study groups for both alpha and beta diversity, the alpha diversity analysis show an intriguing pattern across multiple metrics, which could be validated in future studies with larger cohorts. Specifically, a notable, though non-significant, reduction in alpha diversity was consistently observed in the CeD remission group compared to both active CeD patients and controls across all three metrics evaluated (Shannon diversity index, Chao1 estimator, and Faith phylogenetic diversity; **Figure 2**). This pattern may indicate a link between microbiome dysbiosis and long-term adherence to a GFD as restrictions inherent in the GFD can limit exposure to a broad range of fermentable dietary fibers and other prebiotic compounds that support gut microbial diversity. GFD adherence can also lead to nutritional imbalances, including deficiencies in essential nutrients and total dietary fiber, and may increase exposure to detrimental food components like arsenic, mercury, cadmium, lead, saturated fats, and cholesterol ^25^. These factors, coupled with persistent transcriptomic and metabolic alterations in the intestinal mucosa during disease remission ^26, 27^, including changes in nutrient metabolism ^27^, could contribute to this constrained microbial diversity. Notably, this lower diversity could potentially impact gut health and influence susceptibility of CeD patients to other conditions such as metabolic syndrome, high homocysteine levels, and increased mortality risk due to cardiovascular diseases ^28^.

### Residual dysbiosis persists during CeD remission, characterized by enrichment of inflammation-associated taxa and depletion of protective commensals

Beyond diversity metrics, our DA analyses identified several microbiome alterations consistent with residual dysbiosis in CeD remission. These included enrichment of inflammation-associated taxa and depletion of health-associated commensals, some of which were shared between active CeD and remission. Among remission-specific changes, *Eggerthella lenta* was enriched compared to both controls and active CeD (**Figure 4**; **Supplementary Figure 2A**). *Eggerthella lenta* is a potential pathogenic species that has been found to be elevated in inflammatory conditions such as IBD and rheumatoid arthritis and to activate Th17 cells ^29-31^. We also observed enrichment of *Mogibacterium sp.* NSJ-24 in both CeD remission and active CeD (**Figure 3**; **Supplementary Figure 2C**). The *Mogibacterium* genus, which includes *Mogibacterium* sp. NSJ-24, has been associated with pro-inflammatory responses ^32^. The enrichment of these inflammation-associated taxa suggests that disease-associated microbial features may persist despite mucosal recovery.

Several beneficial species were also depleted in CeD remission relative to either active CeD and/or controls. These species include *Alistipes finegoldii* (**Figure 3; Supplementary Figure 3A**), *Bifidobacterium longum* (**Figures 3** and **4**), *Bacteroides uniformis* (**Figure 4**; non-significant), *Faecalibacterium sp.* (**Figure 3**), *Lacrimispora saccharolytica* (**Figure 3**), *Oscillibacter sp.* MCC667 (**Figure 3**; also see **Supplementary Figure 3A**). Previous studies support the protective roles of these taxa linking them to anti-inflammatory signaling, gut-barrier integrity, and cardiometabolic health. For example, *Alistipes finegoldii* has demonstrated anti-inflammatory properties and enhancement of gut barrier function ^33^, *Bacteroides uniformis* and *Bifidobacterium longum* are well-known probiotics that can inhibit inflammation and regulate the immune system ^34, 35^. *Faecalibacterium* species are consistently reported to be negatively correlated with IBD and colorectal cancer ^36^. *Lachnospira* species have been associated with SCFA production and gut–heart axis modulation ^37, 38^. In particular, *L. saccharolytica* abundance was reported to be significantly reduced in systemic lupus erythematosus ^39^, implying a negative association between this species and autoimmunity. Additionally, *Oscillibacter* species (which include *O. sp.* MCC667) have been linked to cholesterol metabolism and cardiovascular health ^40^.

Additional depletion of potentially protective taxa was shared between active CeD and remission. These included *Anaerostipes hadrus_A* (non-significant), *Dorea sp.* AM13-35 (non-significant), and *CAG*-103 sp000432375 (non-significant) (**Figure 4**; **Supplementary Figure 3C**). Species of *Anaerostipes* have been shown to be involved in butyrate synthesis ^41^, supporting intestinal barrier function and mucosal immunity ^42^. The *Dorea* genera, which includes *Dorea sp.* AM13-35, is part of the *Lachnospiraceae* family, which had been shown to decrease in abundance in IBD patients ^43^. *CAG-103 sp000432375* is a member of the *Oscillospiraceae* family that contribute to the production of valeric acid, which is positively correlated with anti-inflammatory response ^44^.

Together, the enrichment of inflammation-associated taxa and depletion of multiple protective commensals indicate incomplete microbial recovery despite mucosal recovery in CeD remission. The presence of some of these alterations in both active CeD and remission further supports the persistence of disease-associated microbial dysregulation and possible ongoing subclinical inflammation even after histologic recovery.

### CeD remission also exhibits adaptive microbiome remodeling through enrichment of protective taxa and suppression of inflammation-associated microbes

Despite evidence of residual dysbiosis, CeD remission was also characterized by microbial changes that may support mucosal recovery and reduced immune activation. These included enrichment of short-chain fatty acid (SCFA)-producing and potentially protective taxa together with depletion of inflammation-associated microbes. In particular, our DA analyses identified remission-specific enrichment of *Lachnospiraceae bacterium* 210521-DFI.1.109 (compared to both controls and active CeD) and *Ruminococcus lactaris (*compared to controls) (**Figures 3** and **4**; **Supplementary Figure 2A**). Members of the *Lachnospiraceae* family are well-known producers of SCFAs ^37, 38^ as noted before.

Similarly, species of *Ruminococcus* are reported to play a predominant role in the fermentation of dietary fiber to produce SCFA thereby contributing to mucosal recovery and immune regulation ^45^. We additionally observed a non-significant increase in the abundance of three *Blautia* species, namely *Blautia caecimuris*, *Blautia hydrogenotrophica*, and *Blautia schinkii* in CeD remission (**Figure 3**; **Supplementary Figure 2A**). While species of the *Blautia* genera remain relatively understudied, certain *Blautia* species have been reported to have a negative correlation with and reduced abundance in patients with Crohn’s disease ^46^, indicating a potentially protective role.

We further identified depleted taxa in CeD remission that represent suppression of inflammation-associated microbes during remission. These include an uncultured *Dialister* species and *Eggerthellaceae bacterium* (**Figure 3**). Species of *Dialister* are reported to be enriched in Crohn’s disease patients and was associated with increased disease activity ^47^. Species of *Eggerthellaceae* have been also reported to be enriched in several autoimmune diseases, including Sjögren’s syndrome, systemic lupus erythematosus, and multiple sclerosis ^29^. Depletion of these taxa in patients with CeD histologic remission suggests that remission under a GFD may involve selective suppression of inflammation-associated taxa, potentially contributing to reduced immune activation despite persistent microbial imbalance.

Overall, these findings suggest that CeD histologic remission involves adaptive microbiome remodeling characterized by enrichment of SCFA-producing and potentially protective taxa together with selective suppression of inflammation-associated microbes. These coordinated changes may help reduce immune activation, support intestinal barrier function, and promote mucosal recovery during remission.

### Active CeD exhibits dysbiotic microbial disruption characterized by enrichment of inflammation-associated taxa and depletion of protective commensals, alongside potentially compensatory microbial shifts

Active CeD was characterized by a complex pattern of microbial alterations involving enrichment of both inflammation-associated and potentially protective taxa, together with depletion of several SCFA-producing and immunomodulatory commensals. Twelve species were enriched in active CeD relative to CeD remission and/or controls (also **Figures 3** and **4** and **Supplementary Figure 2B**). Of these, *Neobittarella massiliensis* (**Figure 3**) is a recently isolated gut species and little is currently known regarding its functional role. *Bacteroides fragilis* and *Ruminococcus callidus* (**Figure 3**) were previously linked to inflammation or autoimmune pathology ^48-50^, aligning with expectations for an active disease state. The elevated abundance of these species may contribute directly to the inflammatory processes underlying active disease state. In contrast, the other nine species enriched in active CeD—including *Bacteroides xylanisolvens* (**Figure 3**), *Bifidobacterium catenulatum* (**Figure 3**), *Blautia sp.* 210820-DFI.6.14 (**Figure 3**; non-significant), *Coprococcus eutactus_A* (**Figure 4**; non-significant), *Faecalibacterium* sp Marseille-Q4137 (**Figure 3**), *Fimenecus* sp000432435 (**Figure 3**), *Oliverpabstia intestinalis* (**Figure 4**), *Parabacteroides distasonis* (**Figure 3**), *Romboutsia timonensis* (**Figure 4**)—have anti-inflammatory or potential protective effects ^36, 46, 51-59^, suggesting that ongoing inflammation may be accompanied by compensatory microbial restructuring.

We also observed several (potentially) protective species that were depleted in the active CeD relative to healthy controls. These species included *Acetatifactor sp.* (**Figure 3**), *Bifidobacterium sp.* CLA-AA-H280 (**Figure 3**), *Dorea_A longicatena* (**Figure 4**), *Faecalibacterium sp.* CLA-AA-H233 (**Figure 3**), *Gordonibacter urolithinfaciens* (**Figure 3**), *Ruminococcus sp.* AM42-11, and an uncultured species *Eubacteriales* bacterium (**Figures 3**; also see **Supplementary Figure 3B**). Except for *Ruminococcus sp.* AM42-11, which has not been functionally characterized, the rest of these species contribute to SCFA production, gut barrier integrity, and anti-inflammatory signaling based on direct and indirect evidence ^36-38, 60-65^.

Together, the concurrent enrichment of pathogenic and (potentially) commensal species, alongside the depletion of protective commensal taxa, may reflect an unstable gut ecosystem during active disease in which ongoing inflammation drives adaptative microbial restructuring as an attempt to counteract chronic inflammation and restore intestinal homeostasis. At the same time, the depletion of the protective commensal taxa indicates a substantially reduced microbial capacity to support epithelial barrier integrity and immune regulation, reflecting an impaired mucosal protection during active disease. Notably, unlike the remission state—where a pathogenic species was also depleted—no pathogenic taxa were found diminished in active CeD. This highlights the more limited capacity of the active disease microbiome to achieve partial microbiome balance compared to the remission state.

### CeD remission shows functional rewiring toward anti-inflammatory functions at the expense of broader metabolic and detoxification pathways

Beyond taxonomic shifts, CeD remission samples exhibited distinctive alterations in functional pathways. Specifically, propionate (propanoate) metabolism was significantly enriched in remission compared to active CeD while fatty acid biosynthesis showed a marked, albeit not statistically significant, enrichment in CeD remission compared to the other two groups (**Figure 5**). Propionate metabolism, which regulates CD4+ T helper cell activation, has been shown to be downregulated in IBD, indicating its anti-inflammatory potential ^66^, aligning with its enrichment in disease remission in our cohort. Enrichment of fatty acid biosynthesis pathway may indicate enhanced microbial capacity for lipid-derived metabolite production in remission, including SCFAs. These metabolites have been shown to improve clinical and histological outcomes of IBD in a controlled trial setting ^67^ and to accelerate small intestinal healing in gluten-sensitized HLA-DQ8 mice during GFD treatment following dietary fiber supplementation ^18^. These findings support a potential role for microbial SCFA production in mucosal recovery during CeD remission. We additionally identified three pathways that were depleted in CeD remission: pentose and glucuronate interconversions and pyrimidine metabolism that were significantly depleted in remission compared to either active CeD or controls, respectively, and nitrotoluene degradation that exhibits a non-significant depletion in remission (**Figure 5**). These pathways provide protective benefits to the host. Pentose and glucuronate interconversions supply energy via carbohydrate metabolism ^68^, whereas pyrimidine metabolism supports cellular proliferation, differentiation, and apoptosis ^69^. Nitrotoluenes are a group of chemicals that may cause a variety of diseases (e.g. anemia, hypercholesterolemia, testicular atrophy) ^70^. Although studies into the clinical effects of nitrotoluene degradation are limited, one study reported on alterations in cancer genes and proteins that likely contributed to large intestinal tumors in mice following o-nitrotoluene exposure ^71^, implying that a decrease in abundance or activity of nitrotoluene degradation may indicate an impaired ability to clear a harmful chemical.

Functional alterations were also observed in active CeD. For example, benzoate degradation was significantly depleted in active CeD compared with CeD remission (**Figure 5**). The depletion of benzoate degradation may lead to higher benzoate accumulation, promoting secretion of pro-inflammatory cytokines such as IL-1β and IL-6 ^72^, consistent with an active disease state.

Collectively, the enrichment of beneficial pathways modulating immune responses alongside the selective loss of protective pathways during remission reflect major shifts in microbiome’s functional landscape as inflammation subsides and dietary restrictions limit substrate availability. These shifts suggest that, in remission, the microbiome may prioritize anti-inflammatory and mucosal healing functions over broader metabolic activities. This trade-off appears to support remission maintenance but may also constrain the full recovery of metabolic diversity, potentially reducing microbiome resilience and predisposing patients to broader metabolic disturbances.

### Limitations of this study

The principal constraint of this research is the limited size of the study population. This can increase the likelihood of false negative results. Here, we tried to mitigate this limitation by presenting results from two complementary methods for DA analysis (ANCOM-BC and Wilcoxon Rank Sum test), and by reporting non-significant results that show a marked difference between the study groups for descriptive purposes. While this limitation does not affect the reliability of our results when statistical significance is claimed, similar studies with higher sample sizes are needed to confirm non-significant results presented in this study. Nevertheless, although modest by conventional clinical cohort standards, our 21-participant cohort is larger than most published human gut studies employing whole-community PacBio HiFi long-read metagenomics ^73-78^, reflecting in part the historically higher sequencing costs and lower throughput of this approach relative to conventional short-read metagenomics. Another limitation of our study is that all participants receiving a GFD in our cohort were in histologic remission, and no non-responders to the diet (no patients with persistent enteropathy despite a GFD) were included. Consequently, the effects of GFD exposure cannot be statistically disentangled from those of the remission state itself. Furthermore, this study is observational and associative; therefore, the identified microbial alterations cannot be interpreted as causal drivers or consequences of remission. Finally, the generalizability of our findings may be limited by cohort-specific factors such as demographics and dietary patterns, warranting future studies involving more diverse populations from multiple sites to validate and extend our results.

## Conclusions

Overall, our study provides a genome-resolved characterization of the gut microbiota in CeD remission by using long-read metagenomic sequencing. Our findings show that despite mucosal recovery following a GFD, remission is marked by incomplete microbial normalization and distinct functional adaptations, rather than restoration to a healthy baseline. Taxonomically, CeD remission is characterized by potentially adaptive enrichment of SCFA-producing taxa and suppression of inflammation-associated microbes, concurrent with persistent depletion of health-associated commensals and enrichment of some inflammation-associated taxa. Functionally, remission is characterized by enrichment of anti-inflammatory pathways alongside depletion of broader metabolic and detoxification functions. Together, these alterations define a microbial ecosystem distinct from both healthy controls and patients with active disease. Although metabolically constrained, this ecosystem appears to represent a distinct adaptive, but incomplete, state that counterbalances persistent dysbiosis and residual inflammation through protective mechanisms that may limit immune activation, support mucosal healing, and promote immune tolerance. While the limited cohort size, inability to parse the effects of GFD from remission, and observational nature of the study necessitates future validation and larger and more diverse populations to confirm our findings, this study advances our understanding of the gut microbiota’s role and its interactions with the immune and metabolic systems in CeD remission. Collectively, this work lays a strong foundation for subsequent research aimed at unraveling the mechanisms underpinning disease remission and identifying potential therapeutic targets within the gut microbiota.

## Methods

### Study cohort and biospecimens

Stool samples for this study were obtained from an existing biorepository of patients with CeD and controls at Massachusetts General Hospital (MGH), Department of Pediatrics, as part of the Pediatric Biorepository (PBR) study. These biospecimens were obtained from pediatric patients aged 3 to 20 years old. CeD remission was defined by mucosal recovery on follow-up histology (Marsh 0–1) after adherence to a GFD. The PBR study was approved by the Mass General Brigham Institutional Review Board (IRB protocol no. 2016P000949), and informed consent was obtained from participants or their legal guardians, with assent obtained from minors as applicable.

### DNA extraction

Stool samples were collected either with or without RNAlater (Invitrogen™, catalog no. AM7020). For samples preserved in RNAlater, the preservative was removed before DNA extraction. Briefly, cryotubes containing stool samples were thawed on ice for 1 h. A 500-µL aliquot of each sample was transferred to a 1.5-mL microcentrifuge tube and centrifuged at 8,000 rpm for 3 min. The supernatant containing RNAlater was removed, and the resulting pellet was resuspended in 500 µL of 1× phosphate-buffered saline (PBS; Gibco™, pH 7.4, catalog no. 10010023). The samples were centrifuged again at 8,000 rpm for 3 min, and the supernatant was removed to eliminate residual RNAlater. DNA was then extracted from the pellet using the DNeasy PowerSoil Pro Kit (QIAGEN, catalog no. 47014) according to the manufacturer’s instructions. For stool samples collected without RNAlater, the cryotubes were thawed on ice for 1 h. A 0.25-g aliquot of each stool sample was used for DNA extraction with the DNeasy PowerSoil Pro Kit according to the manufacturer’s instructions. DNA concentrations in stool and breast milk samples were measured using the Quant-iT™ PicoGreen™ double-stranded DNA Assay Kit (Invitrogen™, catalog no. P11496) according to the manufacturer’s instructions.

### Long-read HiFi metagenomic sequencing

Extracted DNA underwent PacBio’s long-read HiFi metagenomic sequencing at Maryland Genomics (Institute for Genome Sciences, Baltimore, MD, USA). Sample were sent for sequencing in 96-well plate (BioRad, Hard-Shell® 96-Well PCR Plates, low profile, thin wall, skirted, red/clear #HSP9611). Long-read metagenomic sequencing was performed by using the PacBio Sequel II 8M SMRT Cell run (CCS/HiFi mode – 30-hour movie).

### MAGs reconstruction and taxonomic assignment

MAGs were reconstructed independently for each sample. HiFi reads were first assembled using hifiasm-meta ^79^ with default parameters. The resulting assembly contigs, together with the corresponding HiFi reads, were then processed using the PacBio HiFi-MAG-Pipeline v2.0 ^80^, a completeness-aware binning workflow for long-read metagenomic assemblies. The pipeline uses a completeness-aware binning strategy designed to prevent long, near-complete contigs from being incorrectly grouped with shorter contigs during conventional binning. The strategy identifies all contigs longer than 500 kb and evaluates them individually for genome completeness using CheckM2 ^81^. Contigs estimated to be >93% complete were retained as individual MAGs and excluded from subsequent binning. The remaining incomplete contigs were independently binned using MetaBAT2 ^82^ and SemiBin2 ^83^, after which the resulting bin sets were dereplicated and merged using DAS Tool ^84^. The quality of the merged bins was reassessed using CheckM2, and bins were retained if they had >70% completeness (based on universal single-copy genes, SCGs), <10% contamination, and <20 contigs. These criteria place the MAG quality between the medium and high-quality standards for short-read MAGs proposed by the Genomic Standards Consortium ^85^. These standards do not include an upper limit on the number of contigs per MAG, though contiguity is an important quality to consider. The final MAG set therefore consisted of both long, highly complete contigs identified prior to binning and high-quality bins reconstructed from the remaining incomplete contigs. Taxonomic assignments for the final MAG set were obtained using GTDB-Tk ^86^.

### Taxonomic profiling of HiFi reads

Taxonomic profiling was performed using the PacBio Taxonomic-Profiling-Diamond-Megan workflow, which implements the protein-based MEGAN-LR ^87^ approach (MEGAN-LR-prot) for long-read metagenomic profiling ^88^. For each sample, HiFi reads were aligned against the NCBI non-redundant protein (nr) database using DIAMOND ^89^ in translated-alignment mode (`blastx`) with parameters `-f 101 -F 5000 --range-culling --top 5 -b 12`. The resulting alignments were merged and converted to MEGAN RMA format using MEGAN-LR, which interprets protein alignments across the length of each read to assign taxonomic classifications. The workflow generated both filtered and unfiltered RMA files and summarized read assignments across taxonomic classes. Taxonomic read counts and classifications were subsequently extracted using the MEGAN6 ^90^ command-line utilities.

### Functional profiling of HiFi reads

Functional annotation of HiFi reads was performed using the same DIAMOND/MEGAN-LR protein-alignment framework described above. MEGAN-LR simultaneously assigns functional annotations to genes identified along individual long reads, allowing functional information to be derived from the same protein alignments used for taxonomic classification. Functional assignments were summarized using MEGAN6 across supported annotation systems, including Enzyme Commission (EC), eggNOG, InterPro2GO, and SEED. For downstream pathway-level profiling, the resulting functional annotation data were processed using MEGAN6 ^90^ (MEGAN Community Edition v6.25.10) and mapped to the respective KEGG pathways. Pathway-level read counts were then exported for each sample and used for subsequent downstream analyses.

### Diversity analysis

The Shannon diversity and Chao1 richness estimator were calculated using the functions `diversity` and `estimateR` with the R package `vegan 2.6-6.1`, respectively. The Faith’s phylogenetic diversity was calculated based on weighted UniFrac. For the weighted UniFrac analysis, a taxonomic tree was first generated using the `classification` function from the R package `taxize 0.9.100`. Species names were used as arguments to the function. This function generated the tree through use of the NCBI database. The generated taxonomic tree was then converted to a `phyloseq` object using the R package `phyloseq 1.48.0`. The UniFrac calculations and PERMANOVA tests were performed using the functions `ordinate`, `UniFrac`, and `adonis2` from the `phyloseq` package. The “weighted” parameter in the `ordinate` and `UniFrac` functions was either inputted as “true” to generate a weighted UniFrac analysis or “false” to generate an unweighted UniFrac analysis. The taxonomic tree was used to calculate the Faith’s phylogenetic diversity index using the `estimate_pd` function from the R package `btools 0.0.1`.

For the beta diversity analysis, the Bray-Curtis dissimilarity metric was determined using the `vegdist` function within the R package `vegan 2.6-6.1` and the PERMANOVA test was executed using the `adonis2` function of `vegan` with permutations’ parameter set to 1000. The `vegan`’s `pcoa` function was used for Principal Coordinate Analysis (PCoA).

### Differential abundance analysis

DA analysis was performed using both ANCOM-BC ^23^ and Wilcoxon Rank Sum (Mann-Whitney U) test. ANCOM-BC analysis (implemented using the `ancombc2` function in the `ANCOM-BC 2.6.0` R package). The Wilcoxon test was performed using the `wilcox.test` function of `stats 4.4.0` R package.

#### DA analysis of read alignment-based species and pathways

ANCOM-BC was applied to relative count data (i.e., Total Sum Scaling [TSS]). For the Wilcoxon test, the count data were first normalized using the Trimmed Mean of M-values (TMM) employing the functions `DGEList`, `normLibSizes`, and `cpm` within the R package `edgeR 4.2.1`^91^. This transformation has been shown to reasonably account for differences in sampling fractions across groups^92^.

#### DA analysis of MAG-assigned species

For each MAG-assigned species, ANCOM-BC was applied directly to the average depth of coverage data. For the Wilcoxon Rank Sum test, relative abundance of each MAG-assigned species was calculated by dividing its average depth of coverage by the sum of average depths across all MAG-assigned species within the same sample. Relative abundances were then transformed using the centered log-ratio (CLR) transformation and analyzed by the Wilcoxon Rank Sum test.

#### Integration of DA results and reporting criteria

All the p-values were adjusted for multiple testing by using the `p.adjust` function in R, although significance was claimed based on the raw p-values to safeguard against false negatives due to the low sample size. Microbiome features identified as differentially abundant by either ANCOM-BC or Wilcoxon (p < 0.05) were retained, and the top two non-significant species with the largest fold changes in abundance from each pairwise comparison were additionally reported for exploratory purposes. All raw p-values as well as p-values (adjusted p-values) are provided in **Supplementary File 2**.

## List of abbreviations

CeD: Celiac Disease
GFD: Gluten-free diet

## Declarations

### Ethics approval and consent to participate

This study was conducted in accordance with the Declaration of Helsinki and was approved by the Mass General Brigham Institutional Review Board (IRB protocol no. 2016P000949). Written informed consent was obtained from participants or their legal guardians, with assent obtained from minors as applicable.

### Clinical trial number

Not applicable.

### Availability of data and materials

All processed data generated in this study are provided in the supplementary materials. Raw PacBio HiFi sequencing reads, and reconstructed metagenome-assembled genomes (MAGs) will be deposited in the NCBI Sequence Read Archive (SRA) and GenBank, respectively.

### Author contributions

ARZ conceived and oversaw the study. DL performed all downstream computational analyses. DP conducted taxonomic and functional profiling as well as MAG assembly. TK performed DNA extraction, under the supervision of AMR. MML oversaw sample collection from patients, identified samples for inclusion in the study, and provided critical feedback on the manuscript. AF provided critical feedback on the study and manuscript. ARZ and DL wrote the manuscript. All authors have read and approved the final version of the manuscript.

## Acknowledgements

The authors thank Analeigha Colarusso and Rosiane Lima for assistance with DNA extraction from a subset of samples.

## Supplementary Information

**Supplementary File 1.** Participant metadata and microbiome profiling data, including species abundances, MAG assembly/quality metrics and GTDB-Tk taxonomy, and functional pathway abundances.

**Supplementary File 2.** Differential abundance analysis results for all tested species and pathways, including raw and FDR-adjusted p-values for each feature.

**Supplementary Figure 1.** Non-significant alpha and beta diversity analysis results.

**Supplementary Figure 2.** Species enriched in CeD remission, active CeD, or both.

**Supplementary Figure 3.** Species depleted in CeD remission, active CeD, or both.

## References

1. Singh P, Arora A, Strand TA, Leffler DA, Catassi C, Green PH, Kelly CP, Ahuja V, Makharia GK. Global Prevalence of Celiac Disease: Systematic Review and Meta-analysis. Clin Gastroenterol Hepatol. 2018;16(6):823–36.e2. Epub 20180316. doi: 10.1016/j.cgh.2017.06.037. PubMed PMID: 29551598.

2. Leonard MM, Sapone A, Catassi C, Fasano A. Celiac Disease and Nonceliac Gluten Sensitivity: A Review. Jama. 2017;318(7):647–56. doi: 10.1001/jama.2017.9730. PubMed PMID: 28810029.

3. Ricaño-Ponce I, Wijmenga C, Gutierrez-Achury J. Genetics of celiac disease. Best Pract Res Clin Gastroenterol. 2015;29(3):399–412. Epub 20150508. doi: 10.1016/j.bpg.2015.04.004. PubMed PMID: 26060105.

4. Olshan KL, Leonard MM, Serena G, Zomorrodi AR, Fasano A. Gut Microbiota in Celiac Disease: Microbes, Metabolites, Pathways and Therapeutics. Expert Rev Clin Immunol. 2020. Epub 2020/10/26. doi: 10.1080/1744666X.2021.1840354. PubMed PMID: 33103934.

5. Chibbar R, Dieleman LA. The Gut Microbiota in Celiac Disease and probiotics. Nutrients. 2019;11(10). Epub 2019/10/05. doi: 10.3390/nu11102375. PubMed PMID: 31590358; PMCID: PMC6836185.

6. Pecora F, Persico F, Gismondi P, Fornaroli F, Iuliano S, de’Angelis GL, Esposito S. Gut Microbiota in Celiac Disease: Is There Any Role for Probiotics? Front Immunol. 2020;11:957. Epub 20200515. doi: 10.3389/fimmu.2020.00957. PubMed PMID: 32499787; PMCID: PMC7243837.

7. Luz VCC, Pereira SG. Celiac disease gut microbiome studies in the third millennium: reviewing the findings and gaps of available literature. Front Med Technol. 2024;6:1413637. Epub 20240916. doi: 10.3389/fmedt.2024.1413637. PubMed PMID: 39355139; PMCID: PMC11444026.

8. Galipeau HJ, Hinterleitner R, Leonard MM, Caminero A. Non-Host Factors Influencing Onset and Severity of Celiac Disease. Gastroenterology. 2024;167(1):34–50. Epub 20240128. doi: 10.1053/j.gastro.2024.01.030. PubMed PMID: 38286392; PMCID: PMC11653303.

9. Leonard MM, Valitutti F, Karathia H, Pujolassos M, Kenyon V, Fanelli B, Troisi J, Subramanian P, Camhi S, Colucci A, Serena G, Cucchiara S, Trovato CM, Malamisura B, Francavilla R, Elli L, Hasan NA, Zomorrodi AR, Colwell R, …, de Villsante GC. Microbiome signatures of progression toward celiac disease onset in at-risk children in a longitudinal prospective cohort study. Proceedings of the National Academy of Sciences. 2021;118(29):e2020322118. doi: doi:10.1073/pnas.2020322118.

10. Leonard MM, Karathia H, Pujolassos M, Troisi J, Valitutti F, Subramanian P, Camhi S, Kenyon V, Colucci A, Serena G, Cucchiara S, Montuori M, Malamisura B, Francavilla R, Elli L, Fanelli B, Colwell R, Hasan N, Zomorrodi AR, …, Team C-G. Multi-omics analysis reveals the influence of genetic and environmental risk factors on developing gut microbiota in infants at risk of celiac disease. Microbiome. 2020;8(1):130. Epub 2020/09/11. doi: 10.1186/s40168-020-00906-w. PubMed PMID: 32917289; PMCID: PMC7488762.

11. Rubio-Tapia A, Hill ID, Semrad C, Kelly CP, Greer KB, Limketkai BN, Lebwohl B. American College of Gastroenterology Guidelines Update: Diagnosis and Management of Celiac Disease. Am J Gastroenterol. 2023;118(1):59–76. Epub 20220921. doi: 10.14309/ajg.0000000000002075. PubMed PMID: 36602836.

12. Herrera-deGuise C, Varela E, Sarrabayrouse G, Pozuelo Del Río M, Alonso VR, Sainz NB, Casellas F, Mayorga LF, Manichanh C, Vidaur FA, Guarner F. Gut Microbiota Composition in Long-Remission Ulcerative Colitis is Close to a Healthy Gut Microbiota. Inflamm Bowel Dis. 2023;29(9):1362–9. doi: 10.1093/ibd/izad058. PubMed PMID: 37655859.

13. Rooks MG, Veiga P, Wardwell-Scott LH, Tickle T, Segata N, Michaud M, Gallini CA, Beal C, van Hylckama-Vlieg JE, Ballal SA, Morgan XC, Glickman JN, Gevers D, Huttenhower C, Garrett WS. Gut microbiome composition and function in experimental colitis during active disease and treatment-induced remission. ISME J. 2014;8(7):1403–17. Epub 20140206. doi: 10.1038/ismej.2014.3. PubMed PMID: 24500617; PMCID: PMC4069400.

14. Zafeiropoulou K, Nichols B, Mackinder M, Biskou O, Rizou E, Karanikolou A, Clark C, Buchanan E, Cardigan T, Duncan H, Wands D, Russell J, Hansen R, Russell RK, McGrogan P, Edwards CA, Ijaz UZ, Gerasimidis K. Alterations in Intestinal Microbiota of Children With Celiac Disease at the Time of Diagnosis and on a Gluten-free Diet. Gastroenterology. 2020;159(6):2039–51.e20. Epub 20200810. doi: 10.1053/j.gastro.2020.08.007. PubMed PMID: 32791131; PMCID: PMC7773982.

15. Sample D, Fouhse J, King S, Huynh HQ, Dieleman LA, Willing BP, Turner J. Baseline Fecal Microbiota in Pediatric Patients With Celiac Disease Is Similar to Controls But Dissimilar After 1 Year on the Gluten-Free Diet. JPGN Rep. 2021;2(4):e127. Epub 20211013. doi: 10.1097/PG9.0000000000000127. PubMed PMID: 37206457; PMCID: PMC10191547.

16. Francavilla A, Ferrero G, Pardini B, Tarallo S, Zanatto L, Caviglia GP, Sieri S, Grioni S, Francescato G, Stalla F, Guiotto C, Crocella L, Astegiano M, Bruno M, Calvo PL, Vineis P, Ribaldone DG, Naccarati A. Gluten-free diet affects fecal small non-coding RNA profiles and microbiome composition in celiac disease supporting a host-gut microbiota crosstalk. Gut Microbes. 2023;15(1):2172955. doi: 10.1080/19490976.2023.2172955. PubMed PMID: 36751856; PMCID: PMC9928459.

17. Quagliariello A, Aloisio I, Bozzi Cionci N, Luiselli D, D’Auria G, Martinez-Priego L, Pérez-Villarroya D, Langerholc T, Primec M, Mičetić-Turk D, Di Gioia D. Effect of Bifidobacterium breve on the Intestinal Microbiota of Coeliac Children on a Gluten Free Diet: A Pilot Study. Nutrients. 2016;8(10). Epub 20161022. doi: 10.3390/nu8100660. PubMed PMID: 27782071; PMCID: PMC5084046.

18. Wulczynski M, Constante M, Galipeau HJ, Blom J, Rueda GH, El-Chaar N, Superdock DK, Jiang S, David LA, Murray JA, Surette MG, Armstrong D, Pinto-Sanchez MI, Bercik P, Caminero A, Verdu EF. Small intestinal microbial fiber metabolism dysfunction in celiac disease. Nat Commun. 2026;17(1). Epub 20260331. doi: 10.1038/s41467-026-70644-4. PubMed PMID: 41917007; PMCID: PMC13039995.

19. Rattray N, Kelly P, Farrell G, Russell R, Hansen R, Edwards C, Gillett P, Gerasimidis K. Gluten free diet does not fully restore the faecal metabolome in paediatric coeliac disease. Research Square. 2026. doi: 10.21203/rs.3.rs-10187795/v1.

20. Kim C, Pongpanich M, Porntaveetus T. Unraveling metagenomics through long-read sequencing: a comprehensive review. J Transl Med. 2024;22(1):111. Epub 20240128. doi: 10.1186/s12967-024-04917-1. PubMed PMID: 38282030; PMCID: PMC10823668.

21. Lozupone C, Knight R. UniFrac: a new phylogenetic method for comparing microbial communities. Appl Environ Microbiol. 2005;71(12):8228–35. doi: 10.1128/aem.71.12.8228-8235.2005. PubMed PMID: 16332807; PMCID: PMC1317376.

22. Lozupone CA, Hamady M, Kelley ST, Knight R. Quantitative and qualitative beta diversity measures lead to different insights into factors that structure microbial communities. Appl Environ Microbiol. 2007;73(5):1576–85. Epub 20070112. doi: 10.1128/aem.01996-06. PubMed PMID: 17220268; PMCID: PMC1828774.

23. Lin H, Peddada SD. Analysis of compositions of microbiomes with bias correction. Nat Commun. 2020;11(1):3514. Epub 20200714. doi: 10.1038/s41467-020-17041-7. PubMed PMID: 32665548; PMCID: PMC7360769.

24. Dallah D, Sulieman H, Zaatreh AA, Kamalov F. Empirical Evaluation of the Relative Range for Detecting Outliers. Entropy (Basel). 2025;27(7). Epub 20250707. doi: 10.3390/e27070731. PubMed PMID: 40724447; PMCID: PMC12295245.

25. Hellman R. Gluten Free Diets - A Challenge for the Practicing Physician. Mo Med. 2020;117(2):119–23. PubMed PMID: 32308234; PMCID: PMC7144711.

26. Leonard MM, Bai Y, Serena G, Nickerson KP, Camhi S, Sturgeon C, Yan S, Fiorentino MR, Katz A, Nath B, Richter J, Sleeman M, Gurer C, Fasano A. RNA sequencing of intestinal mucosa reveals novel pathways functionally linked to celiac disease pathogenesis. PLoS One. 2019;14(4):e0215132. Epub 2019/04/18. doi: 10.1371/journal.pone.0215132. PubMed PMID: 30998704; PMCID: PMC6472737.

27. McCreery CV, Alessi D, Mollo K, Fasano A, Zomorrodi AR. Investigating intestinal epithelium metabolic dysfunction in celiac disease using personalized genome-scale models. BMC Med. 2025;23(1):95. Epub 20250221. doi: 10.1186/s12916-025-03854-0. PubMed PMID: 39984962; PMCID: PMC11846356.

28. Kutlu T. Gluten-free diet: is it really always beneficial? Turk Pediatri Ars. 2019;54(2):73–5. Epub 20190711. doi: 10.14744/TurkPediatriArs.2019.82609. PubMed PMID: 31384140; PMCID: PMC6666359.

29. Chang SH, Choi Y. Gut dysbiosis in autoimmune diseases: Association with mortality. Front Cell Infect Microbiol. 2023;13:1157918. Epub 20230331. doi: 10.3389/fcimb.2023.1157918. PubMed PMID: 37065187; PMCID: PMC10102475.

30. Hertz S, Anderson JM, Nielsen HL, Schachtschneider C, McCauley KE, Özçam M, Larsen L, Lynch SV, Nielsen H. Fecal microbiota is associated with extraintestinal manifestations in inflammatory bowel disease. Ann Med. 2024;56(1):2338244. Epub 20240422. doi: 10.1080/07853890.2024.2338244. PubMed PMID: 38648495; PMCID: PMC11036898.

31. Chen J, Wright K, Davis JM, Jeraldo P, Marietta EV, Murray J, Nelson H, Matteson EL, Taneja V. An expansion of rare lineage intestinal microbes characterizes rheumatoid arthritis. Genome Med. 2016;8(1):43. Epub 20160421. doi: 10.1186/s13073-016-0299-7. PubMed PMID: 27102666; PMCID: PMC4840970.

32. Li Q, Pu Y, Lu H, Zhao N, Wang Y, Guo Y, Guo C. Porphyromonas, Treponema, and Mogibacterium promote IL8/IFNγ/TNFα-based pro-inflammation in patients with medication-related osteonecrosis of the jaw. J Oral Microbiol. 2020;13(1):1851112. Epub 20201123. doi: 10.1080/20002297.2020.1851112. PubMed PMID: 33391627; PMCID: PMC7717612.

33. Fischer F, Romero R, Hellhund A, Linne U, Bertrams W, Pinkenburg O, Eldin HS, Binder K, Jacob R, Walker A, Stecher B, Basic M, Luu M, Mahdavi R, Heintz-Buschart A, Visekruna A, Steinhoff U. Dietary cellulose induces anti-inflammatory immunity and transcriptional programs via maturation of the intestinal microbiota. Gut Microbes. 2020;12(1):1–17. doi: 10.1080/19490976.2020.1829962. PubMed PMID: 33079623; PMCID: PMC7583510.

34. Gauffin Cano P, Santacruz A, Moya Á, Sanz Y. Bacteroides uniformis CECT 7771 Ameliorates Metabolic and Immunological Dysfunction in Mice with High-Fat-Diet Induced Obesity. PLOS ONE. 2012;7(7):e41079. doi: 10.1371/journal.pone.0041079.

35. Yao S, Zhao Z, Wang W, Liu X. Bifidobacterium Longum: Protection against Inflammatory Bowel Disease. Journal of Immunology Research. 2021;2021(1):8030297. doi: 10.1155/2021/8030297.

36. Martín R, Rios-Covian D, Huillet E, Auger S, Khazaal S, Bermúdez-Humarán LG, Sokol H, Chatel J-M, Langella P. Faecalibacterium: a bacterial genus with promising human health applications. FEMS Microbiology Reviews. 2023;47(4). doi: 10.1093/femsre/fuad039.

37. Li Z, Zhou E, Liu C, Wicks H, Yildiz S, Razack F, Ying Z, Kooijman S, Koonen DPY, Heijink M, Kostidis S, Giera M, Sanders I, Kuijper EJ, Smits WK, van Dijk KW, Rensen PCN, Wang Y. Dietary butyrate ameliorates metabolic health associated with selective proliferation of gut Lachnospiraceae bacterium 28-4. JCI Insight. 2023;8(4). Epub 20230222. doi: 10.1172/jci.insight.166655. PubMed PMID: 36810253; PMCID: PMC9977501.

38. Chulenbayeva L, Issilbayeva A, Sailybayeva A, Bekbossynova M, Kozhakhmetov S, Kushugulova A. Short-Chain Fatty Acids and Their Metabolic Interactions in Heart Failure. Biomedicines. 2025;13(2). Epub 20250203. doi: 10.3390/biomedicines13020343. PubMed PMID: 40002756; PMCID: PMC11853371.

39. Ahn JS, Lee YB, Han EJ, Choi YJ, Kim DH, Kwok SK, Choi HK, Chung HJ. Identification of specific gut microbes and their therapeutic potential in ameliorating systemic lupus erythematosus in a mouse model. Life Sci. 2025;374:123684. Epub 20250503. doi: 10.1016/j.lfs.2025.123684. PubMed PMID: 40320135.

40. Li C, Stražar M, Mohamed AMT, Pacheco JA, Walker RL, Lebar T, Zhao S, Lockart J, Dame A, Thurimella K, Jeanfavre S, Brown EM, Ang QY, Berdy B, Sergio D, Invernizzi R, Tinoco A, Pishchany G, Vasan RS, …, Xavier RJ. Gut microbiome and metabolome profiling in Framingham heart study reveals cholesterol-metabolizing bacteria. Cell. 2024;187(8):1834–52.e19. doi: 10.1016/j.cell.2024.03.014.

41. Louis P, Duncan SH, McCrae SI, Millar J, Jackson MS, Flint HJ. Restricted distribution of the butyrate kinase pathway among butyrate-producing bacteria from the human colon. J Bacteriol. 2004;186(7):2099–106. doi: 10.1128/jb.186.7.2099-2106.2004. PubMed PMID: 15028695; PMCID: PMC374397.

42. Fu X, Liu Z, Zhu C, Mou H, Kong Q. Nondigestible carbohydrates, butyrate, and butyrate-producing bacteria. Critical Reviews in Food Science and Nutrition. 2019;59(sup1):S130–S52. doi: 10.1080/10408398.2018.1542587.

43. Zhang Z-J, Qu H-L, Zhao N, Wang J, Wang X-Y, Hai R, Li B. Assessment of Causal Direction Between Gut Microbiota and Inflammatory Bowel Disease: A Mendelian Randomization Analysis. Frontiers in Genetics. 2021; Volume 12 - 2021. doi: 10.3389/fgene.2021.631061.

44. Chen YJ, Ho HJ, Tseng CH, Chen YF, Wang ST, Shieh JJ, Wu CY. Short-chain fatty acids ameliorate imiquimod-induced skin thickening and IL-17 levels and alter gut microbiota in mice: a metagenomic association analysis. Sci Rep. 2024;14(1):17495. Epub 20240730. doi: 10.1038/s41598-024-67325-x. PubMed PMID: 39079980; PMCID: PMC11289318.

45. Lakshmanan AP, Al Zaidan S, Bangarusamy DK, Al-Shamari S, Elhag W, Terranegra A. Increased Relative Abundance of Ruminoccocus Is Associated With Reduced Cardiovascular Risk in an Obese Population. Frontiers in Nutrition. 2022; Volume 9 - 2022. doi: 10.3389/fnut.2022.849005.

46. Liu X, Mao B, Gu J, Wu J, Cui S, Wang G, Zhao J, Zhang H, Chen W. Blautia—a new functional genus with potential probiotic properties? Gut Microbes. 2021;13(1):1875796. doi: 10.1080/19490976.2021.1875796.

47. Boronat-Toscano A, Queipo-Ortuño MI, Monfort-Ferré D, Suau R, Vañó-Segarra I, Valldosera G, Cepero C, Astiarraga B, Clua-Ferré L, Plaza-Andrade I, Aranega-Martín L, Cabrinety L, Abadia de Barbarà C, Castellano-Castillo D, Moliné A, Caro A, Domènech E, Sánchez-Herrero JF, Benaiges-Fernandez R, …, Serena C. Dialister-driven succinate accumulation is associated with disease activity and postoperative recurrence in Crohn’s disease. World J Gastroenterol. 2025;31(45):112618. doi: 10.3748/wjg.v31.i45.112618. PubMed PMID: 41378335; PMCID: PMC12687013.

48. Carrow HC, Batachari LE, Chu H. Strain diversity in the microbiome: Lessons from Bacteroides fragilis. PLoS Pathog. 2020;16(12):e1009056. Epub 20201210. doi: 10.1371/journal.ppat.1009056. PubMed PMID: 33301530; PMCID: PMC7728264.

49. Zamani S, Hesam Shariati S, Zali MR, Asadzadeh Aghdaei H, Sarabi Asiabar A, Bokaie S, Nomanpour B, Sechi LA, Feizabadi MM. Detection of enterotoxigenic Bacteroides fragilis in patients with ulcerative colitis. Gut Pathog. 2017;9:53. Epub 20170915. doi: 10.1186/s13099-017-0202-0. PubMed PMID: 28924454; PMCID: PMC5599888.

50. Sankarasubramanian J, Ahmad R, Avuthu N, Singh AB, Guda C. Gut Microbiota and Metabolic Specificity in Ulcerative Colitis and Crohn’s Disease. Front Med (Lausanne). 2020;7:606298. Epub 20201127. doi: 10.3389/fmed.2020.606298. PubMed PMID: 33330572; PMCID: PMC7729129.

51. Despres J, Forano E, Lepercq P, Comtet-Marre S, Jubelin G, Chambon C, Yeoman CJ, Berg Miller ME, Fields CJ, Martens E, Terrapon N, Henrissat B, White BA, Mosoni P. Xylan degradation by the human gut Bacteroides xylanisolvens XB1A(T) involves two distinct gene clusters that are linked at the transcriptional level. BMC Genomics. 2016;17:326. Epub 20160504. doi: 10.1186/s12864-016-2680-8. PubMed PMID: 27142817; PMCID: PMC4855328.

52. Kim H, Jeong Y, Kang S, You HJ, Ji GE. Co-Culture with Bifidobacterium catenulatum Improves the Growth, Gut Colonization, and Butyrate Production of Faecalibacterium prausnitzii: In Vitro and In Vivo Studies. Microorganisms. 2020;8(5). Epub 20200525. doi: 10.3390/microorganisms8050788. PubMed PMID: 32466189; PMCID: PMC7285360.

53. Gavzy SJ, Kensiski A, Lee ZL, Mongodin EF, Ma B, Bromberg JS. Bifidobacterium mechanisms of immune modulation and tolerance. Gut Microbes. 2023;15(2):2291164. Epub 20231206. doi: 10.1080/19490976.2023.2291164. PubMed PMID: 38055306; PMCID: PMC10730214.

54. Yang R, Shan S, Shi J, Li H, An N, Li S, Cui K, Guo H, Li Z. Coprococcus eutactus, a Potent Probiotic, Alleviates Colitis via Acetate-Mediated IgA Response and Microbiota Restoration. Journal of Agricultural and Food Chemistry. 2023;71(7):3273–84. doi: 10.1021/acs.jafc.2c06697.

55. Kverka M, Zakostelska Z, Klimesova K, Sokol D, Hudcovic T, Hrncir T, Rossmann P, Mrazek J, Kopecny J, Verdu EF, Tlaskalova-Hogenova H. Oral administration of Parabacteroides distasonis antigens attenuates experimental murine colitis through modulation of immunity and microbiota composition. Clin Exp Immunol. 2011;163(2):250–9. Epub 20101119. doi: 10.1111/j.1365-2249.2010.04286.x. PubMed PMID: 21087444; PMCID: PMC3043316.

56. Sun H, Guo Y, Wang H, Yin A, Hu J, Yuan T, Zhou S, Xu W, Wei P, Yin S, Liu P, Guo X, Tang Y, Yan Y, Luo Z, Wang M, Liang Q, Wu P, Zhang A, …, Zhou W. Gut commensal Parabacteroides distasonis alleviates inflammatory arthritis. Gut. 2023;72(9):1664–77. Epub 20230105. doi: 10.1136/gutjnl-2022-327756. PubMed PMID: 36604114.

57. Deng J, Qiu Q, Ye S, Yu J, Yao D, Deng H, Wang C, Han L, Deng Y, Chen Y, Liu Y, Liu C, Shang X, Fang X, Lu C. Disentangling environmental and disease-specific signatures in the gut microbiome of psoriasis: discovery of Fimenecus sp. as a novel biomarker and characterization of the gut virome. Journal of Translational Medicine. 2026;24(1):646. doi: 10.1186/s12967-026-08013-4.

58. Wylensek D, Hitch TCA, Riedel T, Afrizal A, Kumar N, Wortmann E, Liu T, Devendran S, Lesker TR, Hernández SB, Heine V, Buhl EM, M. D’Agostino P, Cumbo F, Fischöder T, Wyschkon M, Looft T, Parreira VR, Abt B, …, Clavel T. A collection of bacterial isolates from the pig intestine reveals functional and taxonomic diversity. Nature Communications. 2020;11(1):6389. doi: 10.1038/s41467-020-19929-w.

59. Fan Z, Ke X, Jiang L, Zhang Z, Yi M, Liu Z, Cao J, Lu M, Wang M. Genomic and biochemical analysis reveals fermented product of a putative novel Romboutsia species involves the glycometabolism of tilapia. Aquaculture. 2024;581:740483. doi: 10.1016/j.aquaculture.2023.740483.

60. Xiong Z, Dodson BP, Rogers MB, Sneiderman CT, Janesko-Feldman K, Vagni V, Manole M, Li X, Rajasundaram D, Clark RSB, Raphael I, Morowitz MJ, Mariño E, Kochanek PM, Jha RM, Kohanbash G, Simon DW. Microbial production of short-chain fatty acids attenuates long-term neurologic impairment after traumatic brain injury. Journal of Neuroinflammation. 2025;22(1):285. doi: 10.1186/s12974-025-03615-z.

61. Akobeng AK, Singh P, Kumar M, Al Khodor S. Role of the gut microbiota in the pathogenesis of coeliac disease and potential therapeutic implications. European Journal of Nutrition. 2020;59(8):3369–90. doi: 10.1007/s00394-020-02324-y.

62. Shetty SS, Shetty S, Kumari NS. Therapeutic efficacy of gut microbiota-derived polyphenol metabolite Urolithin A. Beni-Suef University Journal of Basic and Applied Sciences. 2024;13(1):31. doi: 10.1186/s43088-024-00492-y.

63. Mukherjee A, Lordan C, Ross RP, Cotter PD. Gut microbes from the phylogenetically diverse genus Eubacterium and their various contributions to gut health. Gut Microbes. 2020;12(1):1802866. doi: 10.1080/19490976.2020.1802866. PubMed PMID: 32835590; PMCID: PMC7524325.

64. Liang X, Fu Y, Cao WT, Wang Z, Zhang K, Jiang Z, Jia X, Liu CY, Lin HR, Zhong H, Miao Z, Gou W, Shuai M, Huang Y, Chen S, Zhang B, Chen YM, Zheng JS. Gut microbiome, cognitive function and brain structure: a multi-omics integration analysis. Transl Neurodegener. 2022;11(1):49. Epub 20221114. doi: 10.1186/s40035-022-00323-z. PubMed PMID: 36376937; PMCID: PMC9661756.

65. Mondot S, Lepage P, Seksik P, Allez M, Tréton X, Bouhnik Y, Colombel JF, Leclerc M, Pochart P, Doré J, Marteau P. Structural robustness of the gut mucosal microbiota is associated with Crohn’s disease remission after surgery. Gut. 2016;65(6):954–62. Epub 20151201. doi: 10.1136/gutjnl-2015-309184. PubMed PMID: 26628508; PMCID: PMC4893116.

66. Ning L, Zhou Y-L, Sun H, Zhang Y, Shen C, Wang Z, Xuan B, Zhao Y, Ma Y, Yan Y, Tong T, Huang X, Hu M, Zhu X, Ding J, Zhang Y, Cui Z, Fang J-Y, Chen H, Hong J. Microbiome and metabolome features in inflammatory bowel disease via multi-omics integration analyses across cohorts. Nature Communications. 2023;14(1):7135. doi: 10.1038/s41467-023-42788-0.

67. Morrison DJ, Preston T. Formation of short chain fatty acids by the gut microbiota and their impact on human metabolism. Gut Microbes. 2016;7(3):189–200. Epub 20160310. doi: 10.1080/19490976.2015.1134082. PubMed PMID: 26963409; PMCID: PMC4939913.

68. Lin X-L, Guo F, Rillig MC, Chen C, Duan G-L, Zhu Y-G. Effects of common artificial sweeteners at environmentally relevant concentrations on soil springtails and their gut microbiota. Environment International. 2024;185:108496. doi: 10.1016/j.envint.2024.108496.

69. Wang W, Cui J, Ma H, Lu W, Huang J. Targeting Pyrimidine Metabolism in the Era of Precision Cancer Medicine. Frontiers in Oncology. 2021; Volume 11 - 2021. doi: 10.3389/fonc.2021.684961.

70. Deng Y, Meyer SA, Guan X, Escalon BL, Ai J, Wilbanks MS, Welti R, Garcia-Reyero N, Perkins EJ. Analysis of common and specific mechanisms of liver function affected by nitrotoluene compounds. PLoS One. 2011;6(2):e14662. Epub 20110208. doi: 10.1371/journal.pone.0014662. PubMed PMID: 21346803; PMCID: PMC3035612.

71. Sills RC, Hong HL, Flake G, Moomaw C, Clayton N, Boorman GA, Dunnick J, Devereux TR. o-Nitrotoluene-induced large intestinal tumors in B6C3F1 mice model human colon cancer in their molecular pathogenesis. Carcinogenesis. 2004;25(4):605–12. Epub 20031219. doi: 10.1093/carcin/bgh044. PubMed PMID: 14688030.

72. Xiao N, Ruan S, Mo Q, Zhao M, Feng F. The Effect of Sodium Benzoate on Host Health: Insight into Physiological Indexes and Gut Microbiota. Foods. 2023;12(22). Epub 20231110. doi: 10.3390/foods12224081. PubMed PMID: 38002138; PMCID: PMC10670719.

73. Gehrig JL, Portik DM, Driscoll MD, Jackson E, Chakraborty S, Gratalo D, Ashby M, Valladares R. Finding the right fit: evaluation of short-read and long-read sequencing approaches to maximize the utility of clinical microbiome data. Microb Genom. 2022;8(3). doi: 10.1099/mgen.0.000794. PubMed PMID: 35302439; PMCID: PMC9176275.

74. Plaza Oñate F, Roume H, Almeida M. Recovery of Metagenome-Assembled Genomes from a Human Fecal Sample with Pacific Biosciences High-Fidelity Sequencing. Microbiol Resour Announc. 2022;11(6):e0025022. Epub 20220509. doi: 10.1128/mra.00250-22. PubMed PMID: 35532226; PMCID: PMC9202402.

75. Kim CY, Ma J, Lee I. HiFi metagenomic sequencing enables assembly of accurate and complete genomes from human gut microbiota. Nat Commun. 2022;13(1):6367. Epub 20221026. doi: 10.1038/s41467-022-34149-0. PubMed PMID: 36289209; PMCID: PMC9606305.

76. Takewaki D, Kiguchi Y, Masuoka H, Manu MS, Raveney BJE, Narushima S, Kurokawa R, Ogata Y, Kimura Y, Sato N, Ozawa Y, Yagishita S, Araki T, Miyake S, Sato W, Suda W, Yamamura T. Tyzzerella nexilis strains enriched in mobile genetic elements are involved in progressive multiple sclerosis. Cell Rep. 2024;43(10):114785. Epub 20240927. doi: 10.1016/j.celrep.2024.114785. PubMed PMID: 39341204.

77. Minich JJ, Allsing N, Din MO, Tisza MJ, Maleta K, McDonald D, Hartwick N, Mamerto A, Brennan C, Hansen L, Shaffer J, Murray ER, Duong T, Knight R, Stephenson K, Manary MJ, Michael TP. Culture-independent meta-pangenomics enabled by long-read metagenomics reveals associations with pediatric undernutrition. Cell. 2025;188(23):6666–86.e25. Epub 20250909. doi: 10.1016/j.cell.2025.08.020. PubMed PMID: 40930091.

78. Fan Y, Ni M, Aggarwala V, Mead EA, Ksiezarek M, Cao L, Kamm MA, Borody TJ, Paramsothy S, Kaakoush NO, Grinspan A, Faith JJ, Fang G. Long-read metagenomics for strain tracking after faecal microbiota transplant. Nat Microbiol. 2025;10(12):3258–71. Epub 20251022. doi: 10.1038/s41564-025-02164-8. PubMed PMID: 41125958; PMCID: PMC12967305.

79. Feng X, Cheng H, Portik D, Li H. Metagenome assembly of high-fidelity long reads with hifiasm-meta. Nat Methods. 2022;19(6):671–4. Epub 20220509. doi: 10.1038/s41592-022-01478-3. PubMed PMID: 35534630; PMCID: PMC9343089.

80. Portik DM, Feng X, Benoit G, Nasko DJ, Auch B, Bryson SJ, Cano R, Carlin M, Damerum A, Farthing B, Grove JR, Islam M, Langford KW, Liachko I, Locken K, Mangelson H, Tang S, Zhang S, Quince C, Wilkinson JE. Highly accurate metagenome-assembled genomes from human gut microbiota using long-read assembly, binning, and consolidation methods. bioRxiv. 2024:2024.05.10.593587. doi: 10.1101/2024.05.10.593587.

81. Chklovski A, Parks DH, Woodcroft BJ, Tyson GW. CheckM2: a rapid, scalable and accurate tool for assessing microbial genome quality using machine learning. Nat Methods. 2023;20(8):1203–12. Epub 20230727. doi: 10.1038/s41592-023-01940-w. PubMed PMID: 37500759.

82. Kang DD, Li F, Kirton E, Thomas A, Egan R, An H, Wang Z. MetaBAT 2: an adaptive binning algorithm for robust and efficient genome reconstruction from metagenome assemblies. PeerJ. 2019;7:e7359. Epub 20190726. doi: 10.7717/peerj.7359. PubMed PMID: 31388474; PMCID: PMC6662567.

83. Pan S, Zhao XM, Coelho LP. SemiBin2: self-supervised contrastive learning leads to better MAGs for short- and long-read sequencing. Bioinformatics. 2023;39(39 Suppl 1):i21-i9. doi: 10.1093/bioinformatics/btad209. PubMed PMID: 37387171; PMCID: PMC10311329.

84. Sieber CMK, Probst AJ, Sharrar A, Thomas BC, Hess M, Tringe SG, Banfield JF. Recovery of genomes from metagenomes via a dereplication, aggregation and scoring strategy. Nat Microbiol. 2018;3(7):836–43. Epub 20180528. doi: 10.1038/s41564-018-0171-1. PubMed PMID: 29807988; PMCID: PMC6786971.

85. Bowers RM, Kyrpides NC, Stepanauskas R, Harmon-Smith M, Doud D, Reddy TBK, Schulz F, Jarett J, Rivers AR, Eloe-Fadrosh EA, Tringe SG, Ivanova NN, Copeland A, Clum A, Becraft ED, Malmstrom RR, Birren B, Podar M, Bork P, …, Consortium GS. Minimum information about a single amplified genome (MISAG) and a metagenome-assembled genome (MIMAG) of bacteria and archaea. Nat Biotechnol. 2017;35(8):725–31. doi: 10.1038/nbt.3893. PubMed PMID: 28787424; PMCID: PMC6436528.

86. Chaumeil PA, Mussig AJ, Hugenholtz P, Parks DH. GTDB-Tk: a toolkit to classify genomes with the Genome Taxonomy Database. Bioinformatics. 2019;36(6):1925–7. Epub 20191115. doi: 10.1093/bioinformatics/btz848. PubMed PMID: 31730192; PMCID: PMC7703759.

87. Huson DH, Albrecht B, Bağcı C, Bessarab I, Górska A, Jolic D, Williams RBH. MEGAN-LR: new algorithms allow accurate binning and easy interactive exploration of metagenomic long reads and contigs. Biol Direct. 2018;13(1):6. Epub 20180420. doi: 10.1186/s13062-018-0208-7. PubMed PMID: 29678199; PMCID: PMC5910613.

88. Portik DM, Brown CT, Pierce-Ward NT. Evaluation of taxonomic classification and profiling methods for long-read shotgun metagenomic sequencing datasets. BMC Bioinformatics. 2022;23(1):541. Epub 20221213. doi: 10.1186/s12859-022-05103-0. PubMed PMID: 36513983; PMCID: PMC9749362.

89. Buchfink B, Reuter K, Drost HG. Sensitive protein alignments at tree-of-life scale using DIAMOND. Nat Methods. 2021;18(4):366–8. Epub 20210407. doi: 10.1038/s41592-021-01101-x. PubMed PMID: 33828273; PMCID: PMC8026399.

90. Bağcı C, Beier S, Górska A, Huson DH. Introduction to the Analysis of Environmental Sequences: Metagenomics with MEGAN. Methods Mol Biol. 2019;1910:591–604. doi: 10.1007/978-1-4939-9074-0_19. PubMed PMID: 31278678.

91. Robinson MD, McCarthy DJ, Smyth GK. edgeR: a Bioconductor package for differential expression analysis of digital gene expression data. Bioinformatics. 2010;26(1):139–40. Epub 20091111. doi: 10.1093/bioinformatics/btp616. PubMed PMID: 19910308; PMCID: PMC2796818.

92. Lin H, Peddada SD. Analysis of microbial compositions: a review of normalization and differential abundance analysis. NPJ Biofilms Microbiomes. 2020;6(1):60. Epub 20201202. doi: 10.1038/s41522-020-00160-w. PubMed PMID: 33268781; PMCID: PMC7710733.

